## Supplemental Figure Table for "Covariance predicts conserved protein residue interactions important to the emergence and continued evolution of SARS-CoV-2 as a human pathogen"

### Supplemental Figure Legend

**Supplemental File S1.** Genomes and accession numbers of β-coronaviruses used in both phylogenic and pan covariance analysis.

**Supplemental File S2.** Tables of covariant pairs identified in both pan and clinical covariance analyses. Raw tables are provided to show purity and percentage for each pair. For each pair in the pan-covariance the number and identities of different residue are shown. Residue position by alignment consensus and the estimated corresponding position in SARS-CoV-2 are both tabulated. "Z" Residue indicate deletions.

**Supplemental File S3**. Table of clusters, residues, and respective genome names for pan covariance analysis. Residue numbering is indicated by SARS-CoV-2. "-" indicates corresponding residues absent in SARS-CoV-2 based on alignment and "Z" indicates deletion.  Calculated ATDS for each cluster is tabulated and provided.

**Supplemental File S4.** List and description of both sets of 126,051 sequences extracted from GISAID used for clinical covariance analysis. Source data for each sequence is provided as a reference.

**Supplemental File S5.** Calculated molecular interactions in the 7JJI.PDB file using Arpeggio. Chain, Residue, and Atom interactions are indicated. All direct and indirect positions are indicated. All pan covariant pairs for Spike are listed by abundance of independently identified amino acids and co-occurrence of indirect and direct interactions predicted by Arpeggio are indicated. Accounting of co-occurrence between pan-covariance independent pair representation and predicted interaction in 7JJI.PDB are tabulated.

**Supplemental File S6.** 538 pairs identified that overlap between pan and clinical analyses. Count of each residue in the pan CoV analysis is presented. Presence in the selected list of dominant circulating variants is indicated for each pair. WHO label for each variant is used for reference.

**Supplemental File S7.** Gephi force mapping file of residues, clusters, CoVs from pan-CoV analysis.

**Supplemental File S8.** Newick file for ML tree generated using 30,533 nucleotide alignment of 169 CoVs between SARS-CoV-2 *Nsp1* start site and *N* stop codon.
